# *Mc*LTP1, a lipid transfer protein isolated from noni seeds induces effective healing of superficial burns

**DOI:** 10.1101/2023.02.04.527120

**Authors:** Bianca Moreira Kurita, Gisele de Fátima Pinheiro Rangel, Liviane Maria Alves Rabelo, Tamiris de Fátima Goebel de Sousa, Fernanda Soares Macêdo, Renata Ferreira de Carvalho Leitão, Hermógenes David de Oliveira, Nylane Maria Nunes de Alencar

**Author notes:** **Corresponding author**, **Corresponding author: Prof. Nylane Maria Nunes de Alencar.** Department of Physiology and Pharmacology, Federal University of Ceará, Campus do Porangabuçu, 60430-275, Fortaleza, Ceará, Brazil.

## Abstract

Burns are health problems that overwhelm the Unified Health System (SUS) in Brazil. Despite the new therapeutic strategies, the costs of treating burns ate still quite high, and there are no effective alternatives for healing the skin. The use of plants with therapeutic potential is popularly used, due to its low cost, easy access and great Brazilian biodiversity. *Mc*LTP1, a lipid transfer protein isolated from *Morinda citrifollia* (noni) seeds, has shown antinociceptive, anti-inflammatory, antibacterial and antioxidative effects. Therefore, the aim of this study was to investigate the effect of McLTP1 on the healing of superficial burns in mice. The study was approved by CEUA NPDM – UFC (protocol: 02170619-0). The burn was induced by direct contact with a square stainless-steel plate (1.5 cm^2^). The animals were divided into five experimental groups (n=6-7/grupo) and treated daily with 0.9% NaCl saline solution (Sham), or with topical treatment performed with dermatological creams: Silver sulfadiazine 1% (Sulfa 1%), lanette cream (Vehicle), cream lanette containing 0.25% and 0.5% of *Mc*LTP1. The animals were euthanized after 14 days. *Mc*LTP1 promoted total wound closure after 2 weeks of treatment, reduced histopathological scores at 3^rd^ day, as well as induced the formation of a thicker epithelium and collagens synthesis on 14^th^ day, modulated inflammation by reducing MPO activity, TNF-α, IL-1β and IL-6 levels and increasing IL-10 after 3 days of burn, modulated VEGF production at three times analyzed in this study, increased TGF-β and immunostaining for FGF after 7 days, reduced immunostaining for TNF-α on the 3^rd^ day and exerted an antioxidant function by reducing MDA and nitrite and increasing GSH at day 3. In short, *Mc*LTP1 showed an important healing action in this burn model, showing additional anti-inflammatory and antioxidant effects.

## Introduction

Around the world, around 180,000 patients die from burns each year, predominantly in poor countries^1^. Burns are considered a health problem, since they can be highly aggressive depending on their extension^2^. Enhanced mortality of burns can be related to neglect of treatments and absence of prevention strategies^3^. They can be classified in stages ranging from I to IV. Stage I consist in epithelial burns, stage II in epithelial and dermal burns, stage III in epithelial, dermal, and hypodermal burns and stage IV aggregates burns that cross the hypodermis and reach muscles, bones, and tendons^4,5^.

Cellular mechanisms of skin burns involve capillary thrombosis, inflammatory mediators, and pro-apoptotic factor product, with nuclear factor kB activation, responsible to induce TNF-α, IL-1β, IL-6 and PGE_2_ release^2,6,7^. Burns also result in increased levels of reactive oxygen species, as superoxide anion, hydrogen peroxide and nitric oxid^8^.

Burn wound healing occurs via cellular proliferation. VEGF induces angiogenesis and TGF-β promotes wound contraction^9,10,11^. Control of secondary infections of burns also plays a key part on burn wound healing, since loss of stratified layer contributes to the entry of skin colonizing microorganisms^12^. To avoid this complications, wound cleaning and topic application of antimicrobials agents as bacitracine, nistatine and silver sulfadiazine are employed^13,14^. However, these topic preparations do not have healing effects^15^. In this context, many studies have been investigating the use of medicinal plants as healing agents, not only for burns, but wound in general^16,17,18^.

*Morinda citrifolia L*. (Noni) is a natural plant which medicinal effects haven been related for 2000 years^19,20^. Previous studies founded wide range of therapeutic effects of many parts of Noni, such as antitumoral effects with aqueous extract of leaves and Noni juice^21,22^, antidiabetic effects with Noni juice and root methanolic extract^23,24,25^, antiallergic effects with ethanolic extract of leaves and fruits^26^ (and gastroprotective effects with aqueous extract of fruit^27^. Healing potential of noni was also investigated in excisional wound model in rats, which demonstrated more effective wound contraction after 5 and 11 days^28^.

*Mc*LTP1 is a lipid transfer protein isolated from Noni seeds that has been demonstrating anti-inflammatory, antinociceptive and antioxidant effects^29,30,31,32^. Recently, this protein showed an *in vitro* antibacterial effect in *Sthaphylococcus aureus* and *Staphylococcus epidermidis^33^*, predominant bacteria in secondary skin infections^34,35^.

Considering the promising effects of McLTP1 and the absence of therapeutic alternatives that induce healing effect of skin lesions, it is important to investigate the effect of this protein on the healing of burns.

## Material and Methods

### McLTP1 Isolation and purification

Isolation and purification of *Mc*LTP1 was made in accord with previous studies^29,33^. Noni (Morinda citrifolia L.) seeds were finely ground and deffated with petroleum ether (1:10 w/v), air-dried for 24 h at 25 °C and stored at -20 °C. Purity analysis was made on lyophilized samples of *Mc*LTP1 when submitted do electrophoresis under non-reducing conditions (SDS-PAGE). The total protein concentration was measured by the Bradford method.

### Animals and experimental groups

Adult female Swiss mice *(Mus musculus)*, weighing 25 – 30 g were obtained from the Central Animal House of the Federal University of Ceará and housed on animal house of Drug development research center (24 ± 2°C, 12-h light/dark cycle) with *ad libitum* food and water. The mice were divided in groups of 6–7 animals each. All experimental procedures were conducted following the National Guidelines for the Use of Experimental Animals of Brazil and were approved by the Committee on Ethics in the Use of Animals of the Federal University of Ceará (protocol no. 02170619-0). The mice were randomly divided on five groups that were submitted to superficial burns. One group consisted of non-treated animals (Sham) and the others received silver sulfadiazine 1%, vehicle (lanette) and *Mc*LTP1 at 0,25% e 0,5%.

### Superficial burns induction model

Superficial burns were made in animals with shaved and sanitized (70° ethyl alcohol) backs previously anesthetized with ketamine and xylazine. Burn injury was induced by contact with a stainless-steel hot plate (100 °C – 1.5 cm^2^) for 6 seconds. To avoid secondary infections of the burn wounds, prior to the induction of burns, the benches were cleaned (70° ethyl alcohol). The animals were organized in clean boxes (N=1/cage), with sterile shavings, during the entire duration of the experimental protocol, so that there was no contamination of the burn area by contact with saliva from other animals.

### Wound contraction analysis

The wound areas were first measured on days 0, 3, 5, 7, 9, 12 and 14 to make a temporal course assessment of burn wound healing. This analyze was made in according to previous studies^18^. The areas were taken by using a digital pachymeter and multiplying base x height. Contraction taxes were measured by comparing the areas of 3^rd^, 5^th^, 7^th^, 9^th^, 12^th^ and 14^th^ day to initial area.

### Obtaining tissue samples

After carrying out the time course of the macroscopic evaluation, the 3^rd^, 7^th^ and 14^th^ days were chosen for histological and immunohistochemical evaluation, as well as the mediators involved in inflammation and healing. Euthanasia was performed by anesthetic overdose equivalent to three times the recommended dose for anesthesia in mice, with xylazine hydrochloride (30 mg/kg, i.p.) and ketamine hydrochloride (300 mg/kg, i.p.). The skin on the back of the animals was removed in such a way as to cover the entire extent of the lesions as well as part of the surrounding healthy skin, until the muscle layer was exposed.

### Histologic processing and analysis

The tissue samples were fixed in a 10% (v/v) formaldehyde buffered solution (pH 7.4), replaced by 70% ethyl alcohol after 24 hours and followed for histological processing for inclusion in paraffin at 58°C in the automatic tissue processor (Lupe®). 4 μm-thick segments of the fragments were prepared in a semiautomatic microtome (Leica®) and organized on microscopic slides. The histological evaluation by scores was performed at the experimental times of 3 and 7 days, based on the presence or absence of an ulcer, evaluation of the thickness of the epidermis and dermis, and the presence of inflammatory infiltrate in the dermis and hypodermis^17^, as observed in table 2.

**Table 2.** Histologic analysis scores

| Scores | Parameters |
| --- | --- |
| 0 | No ulcer + epithelium and dermis in normal or restored thickness + absent inflammatory infiltrate + keratinization |
| 1 | Punctual ulcer areas + slight thinning of the epithelium and dermis + discrete inflammatory infiltrate in dermis and hypodermis |
| 2 | Moderate ulcer areas + moderate thinning of the epithelium and dermis + moderate inflammatory infiltrate in the dermis and hypodermis |
| 3 | Extensive ulcer area + loss of epithelium and part of the dermis+ intense inflammatory infiltrate in the dermis and hypodermis |

Recovered epithelium thickness was measured 14 days after injury induction. Using a millimeter ocular lens, the epithelium thickness was measured in five fields of the same slide (same animal) at 100x magnification^36^. After that, the final thickness measurement was the arithmetic mean of the five measurements.

### Myeloperoxidase (MPO) activity dosage

To verify the presence of neutrophils in burnt skins myeloperoxidase activity was evaluated, since this enzyme is abundantly found at neutrophil granules. The samples were removed and stored at -80 °C. On the test day, they were homogenized with buffer solution [NaCl (100 mM), EDTA (15 mM) e NaPO4 (20 mM)] and centrifuged. The pellet was resuspended in buffer solution [NaPO4 (50 mM) and hexadecyl-trimethyl-ammonium bromide] and centrifuged. The supernatant was used for the assay with tetramethylbenzidine (1.6 mM) and hydrogen peroxide (0.5 mM). The enzymatic reaction was stopped with 50 μl of H_2_SO_4_ (1M) in each well. Myeloperoxidase activity was determined at absorbance of 450 nm in a plate reader. The values were expressed as myeloperoxidase activity/mg of tissue^37^.

### Determining TNF-α, IL-1β, IL-6, IL-10, VEGF and TGF-β in burnt skins

Samples collected on the 3rd day were used for TNF-α, IL-1β, IL-6, IL-10 and VEGF dosages. On the 7th and 14th days, the samples were dosed for TGF-β and VEGF. A microtiter plates with 96 well were incubated with anti-TNF-α or anti-IL-1β primary antibodies (R&D Systems) diluted in PBS (1:1000), followed by incubation with biotinylated monoclonal anti-TNF-α, anti-IL-1β, anti-IL6, anti-IL10, anti-VEGF and anti-TGF-β detection antibodies (R&D Systems) diluted in BSA 1% (1:1000). Plates were washed and HRP-streptavidin 1000 (R&D Systems) diluted and were added in each well. Color reagent o-phenylenediamine (R&D Systems) was added and plates were incubated in the absence of light. The enzymatic reactions were stopped with 100 μl of stop solution (2 NH_2_SO_4_). Levels the proteins were determined at absorbance of 450 nm in a plate reader The values were expressed in picograms of cytokines/mg of tissue.

### Immunohistochemistry for TNF-α and FGF

Detection of TNF-α was made at the 3^rd^ day and FGF was assessed at 7^th^ and 14^th^ days. Skin samples were fixed in 10% buffered formalin for 24 hours and processed for inclusion in paraffin for obtention of cuts with a thickness of 4 μm, which were subsequently inserted into silanized histological slides. The slides containing the sections were subjected to deparaffinization in an oven at 60°C for 1 hour, followed by two baths in xylene (5 min). They were hydrated with two baths of absolute ethanol, one bath in 90% ethanol, and one bath in 70% ethanol (3 min). Then, the sections were submerged in a distilled water bath for 10 min and antigenic recovery was performed with citrate buffer (DAKO, pH 6.0) for 25 min in a water bath at a temperature of 95°C. The tissues were washed with phosphate-buffered saline (PBS) for 5 min. In the next step, the peroxidase was blocked with 3% hydrogen peroxide (DAKO) for 30 min. The slides were washed with PBS and incubated with primary anti-TNF-α (AB6671) and anti-FGF (SAB2108135) (1:300 dilution) antibodies overnight. For the preparation of negative controls, the primary antibodies were omitted. After this period, the sections were washed three times with PBS and incubated with HRP polymer (DAKO) for 30 min. Then, the slides were washed three times with PBS for 3 min each, dried and DAB applied (DAKO, 3,3-diaminobenzidine, one drop of DAB for one mL of diluent). The slides were observed until a brown color appeared, after which the reaction was stopped by distilled water. Finally, the slides were counterstained with Harrys hematoxylin and processed to insert the coverslip. Tissue images were captured using a digitized camera attached to the microscope (Nikon Elipse E200), capturing 5 fields per histological section at 400x magnification. The number of marks was counted in each field, and the arithmetic mean per field was calculated for each animal.

### Assessing oxidative stress induced by superficial burns

#### Malondialdehyde levels

Skin samples were collected on the 3rd day and destined for the quantification of malondialdehyde (MDA) levels. MDA is a product of the decomposition of hydroperoxides of polyunsaturated fatty acids, formed during the oxidative process^38^. The reaction involves 2-thiobarbituric acid with MDA, producing a red colored compound, measured spectrophotometrically at 532 nm wavelength. Skin samples were homogenized at 10% (weight/volume) in Politron® with 0.05 M phosphate buffer (pH 7.4). Then, 250 μL of the homogenate were placed in a water bath at 37°C for 1 hour. After this time, in order to interrupt peroxidation, 400 μL of 35% perchloric acid were added to the samples, which were then centrifuged (14000 rpm, 15 minutes, 4°C). From the supernatant obtained, 600 μL were transferred to a microtube, to which 200 μL of 0.8% thiobarbituric acid were added. This mixture was placed in a water bath at 95°C for 30 minutes. After cooling, the samples were plated and the absorbance reading was performed in a microplate reader at a wavelength of 532 nm. A standard curve with known concentrations of tetramethoxypropane (TMP) was used to calibrate the method and the MDA concentration in the samples was calculated using the equation of the straight line for the standard curve. Results were expressed in nmol of MDA/g of tissue.

#### Glutathione reduced levels

Skin samples were collected on the 3rd day and destined for the quantification of reduced glutathione (GSH) levels. GSH is a water-soluble antioxidant recognized as the most important endogenous component of the pool of non-protein sulfhydryl groups (NPSH) in our body. To determine the concentration of GSH, the NP-SH content was analyzed using the technique described by Sedlak and Lindsay (1968), which is based on the reaction of 5,5-dithiobis(2-nitrobenzoic) acid (DTNB) with compounds of sulfhydryl, and consequent development of yellow coloration. DTNB reacts with GSH forming 2-nitro-5-thiobenzoic acid and oxidized glutathione (GSSG). The skin samples were homogenized at 10% (weight/volume) in Politron® with 0.02 M EDTA solution. Soon after, 60 μL of 10% trichloroacetic acid (TCA) were added to 40 μL of each sample in order to to precipitate the proteins present in the biological material. The material was then centrifuged (5000 rpm, 15min, 4°C) and 60 μL of the obtained supernatant was plated. A standard curve with known GSH concentrations was used for method calibration. 102 μL of the reading solution (Tris-EDTA, DTNB 0.01 M) were added and the absorbance was immediately measured in a microplate reader at a wavelength of 412 nm. The GSH concentration in the samples was calculated using the straight-line equation for the standard curve and the results were expressed in μg of GSH/g of tissue.

#### NO conversion to NO2 and NO3 analysis

The skin samples of 3rd day were weighed and crushed (POLITRON®) with the aid of a cold solution of potassium chloride (1.15% KCl), after obtaining the homogenates of each sample (10% tissue), centrifugation was performed to obtaining the supernatant (1500g; 15 minutes). So that all the nitrate (NO^3−^) present in the supernatant was converted into nitrite (NO^2−^), the samples obtained were plated (96-well plate) in duplicate (80 μL of each supernatant) and incubated for 12 hours with a reagent solution (40μL of nitrate reductase enzyme, NADPH substrate, KH_2_PO_4_ in distilled water). A nitrite reference standard curve was also plated from a continuous dilution (1:2) of a 200 μM sodium nitrate (NaNO_2_) solution. After converting nitrate to nitrite, 80 μL of Griess solution (1% suffanilamide in 1% H_3_PO_4_/0.1% NEED/distilled water/1:1:1:1) was added to each well. The reading of the final purple color obtained by the reaction was performed at the absorbance of 540 nm and the results were expressed in μM of NO^2−^, using the standard nitrite curve as a reference to obtain the values.

### Statistical analysis

Data normality was checked by Shapiro-Wilke test. The parametric data were presented as means ± standard error mean (SEM) followed by T test or Univariate Analysis of Variance (ANOVA) and Bonferroni test. The nonparametric data were presented as medians followed by extreme values and analyzed by Mann-Whitney or Kruskal-Wallis test and Dunn post-test. P<0.05 was considered significant. All statistic calculations were performed using GraphPad Prism®, version 8.0 for Mac OS X (La Jolla-CA, EUA), with duly authorized license.

## Results

### Topical treatment with McLTP1 enhances the contraction of burn-induced wounds

As shown in figure 1A, of the dermatological creams used for the treatment of wounds induced by burns, only the one that had McLTP1 as an active ingredient in the highest concentration (0.5%) was efficient in increasing wound contraction throughout the time course of healing in the model used here. McLTP1 at the lowest tested dose (0.25%) did not increase wound contraction at any of the evaluated times. On the other hand, the cream containing 1% silver sulfadiazine, the topical treatment most used in clinical practice for burns, was effective in increasing the rate of wound contraction only on days 3, 5 and 7 after and with a lower percentage of contraction on days 3 and 5, as demonstrated by the cream with McLTP1 (0.5%). Throughout the time course of the evaluation, no increase in the contraction of the lesions was observed in the group of animals treated with the lanette cream without any active principle (vehicle), which rules out the per se effect of the vehicle used in the preparation of the creams. To complement the temporal evaluation of the lesions, they were photographed, with one wound per experimental group illustrated in a representative way of its evolution during the healing process (Figure 1B).

**Figure 1A.**
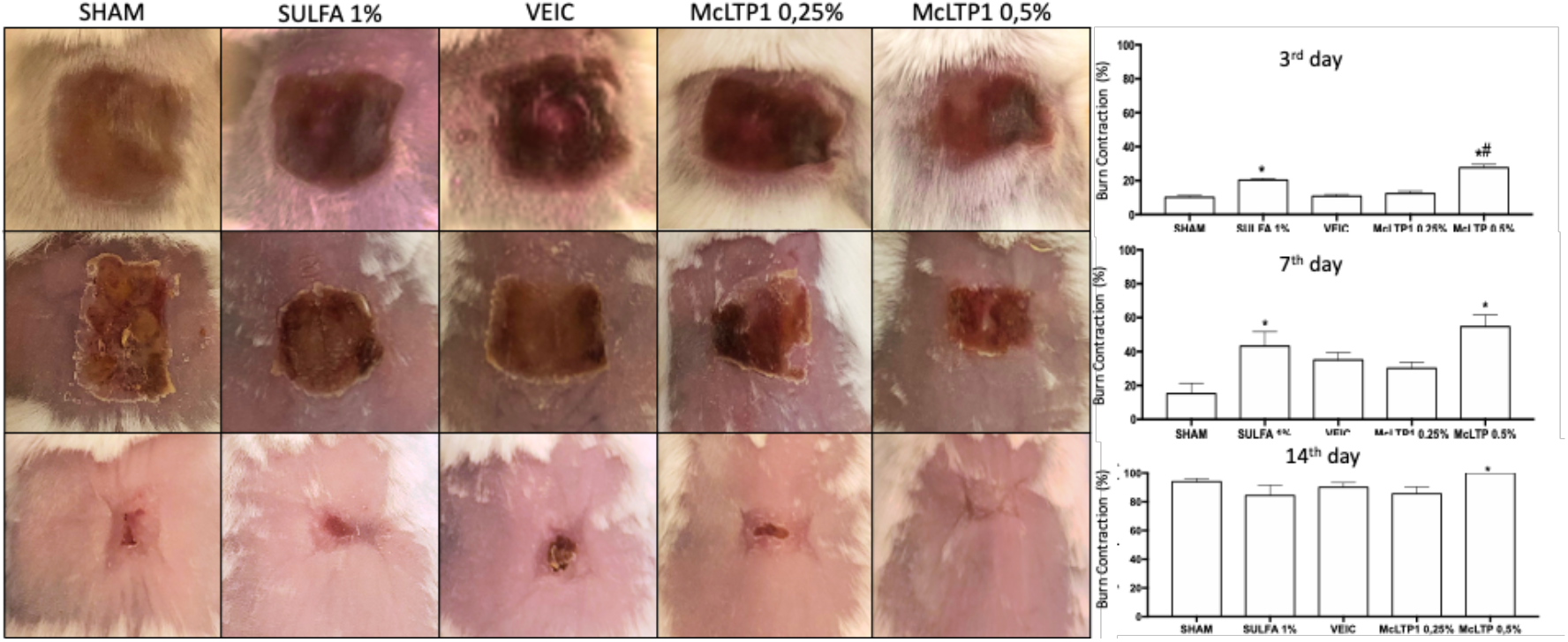
Measurements of wound areas were performed on days 0, 3, 5, 7, 9, 12 and 14 and expressed in percentage of contraction. The animals were treated with a single daily application of dermatological creams for 14 days. McLTP1 (protein at concentrations of 0.25% and 0.5%), Sulfa 1% (silver sulfadiazine 1%), Veic (lanette without addition of active substance) and SHAM (without treatment). ANOVA and Tukey’s post-test were used for comparisons between means. *p<0.05 represents a statistically significant difference in relation to the Sham group, #p<0.05 in relation to Sulfa 1%, δp<0.05 in relation to Vehicle and fp<0.05 in relation to McLTP1 0.25% (N=6-7 animals/group).

**Figure 1B.**
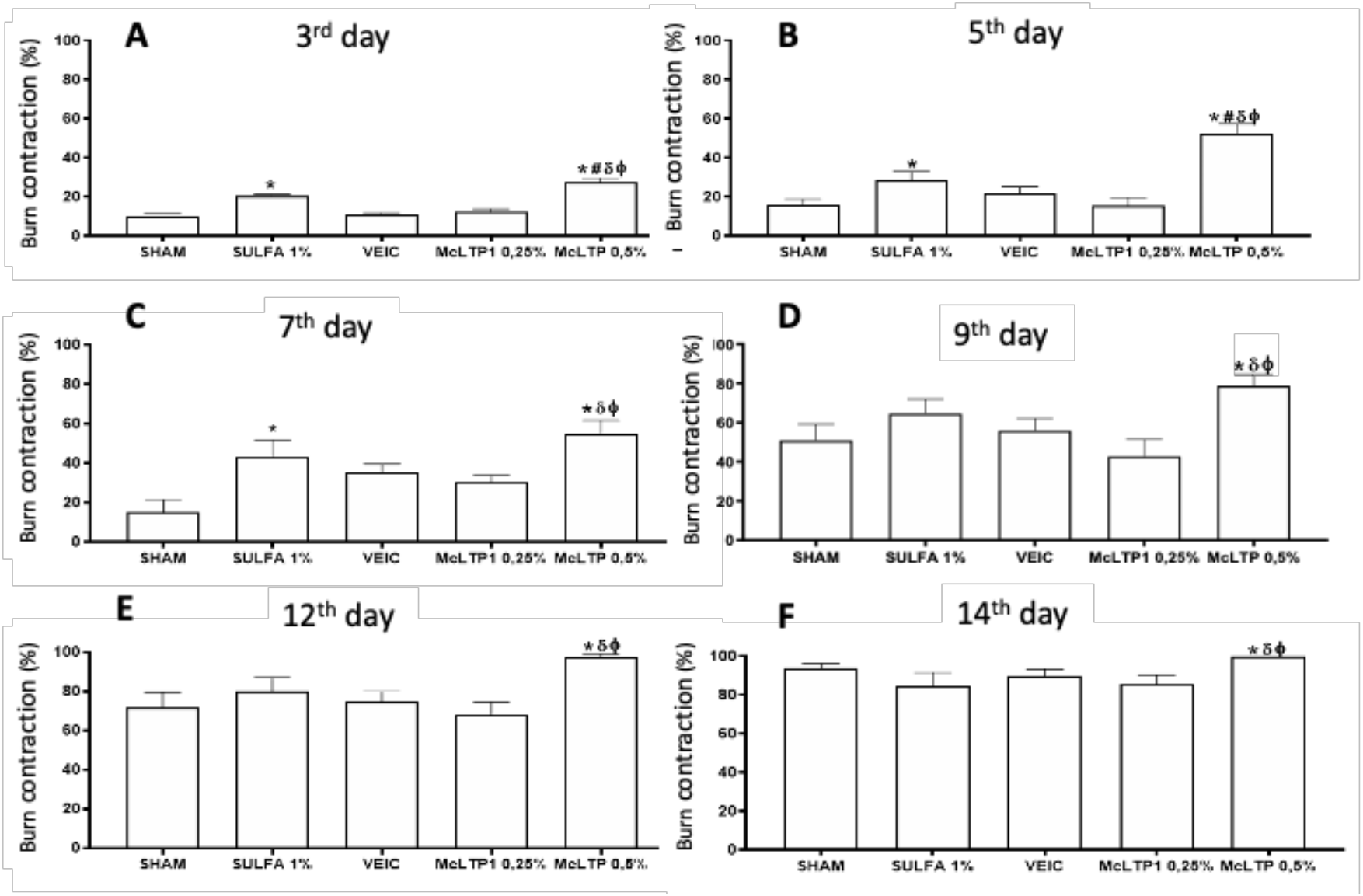
Lesions were photographed on days 3, 7 and 14 after burn induction. One animal per group was chosen to represent the group according to the results obtained by analyzing the contraction of the lesion. The animals were treated with a single daily application of dermatological creams based on McLTP1 (MC) 0.25% and 0.5%, 1% silver sulfadiazine and Vehicle (without active ingredient) for 14 days. (N=6-7 animals/group).

### McLTP1 reduced inflammatory infiltrate and promoted tissue remodeling in burn-induced injuries

On the 3rd day, the histological evaluation by scores showed that the model was efficient in inducing ulcer formation, reducing the thickness of the epithelium and dermis, as well as inducing the presence of inflammatory cells such as polymorphonuclear and mononuclear cells in the dermis and hypodermis. These findings were not observed when evaluating slides from animals not submitted to burn (data not shown). The scores on that day showed no statistical difference between the groups (Chart 1,Figure 2).

**Figure 2.**
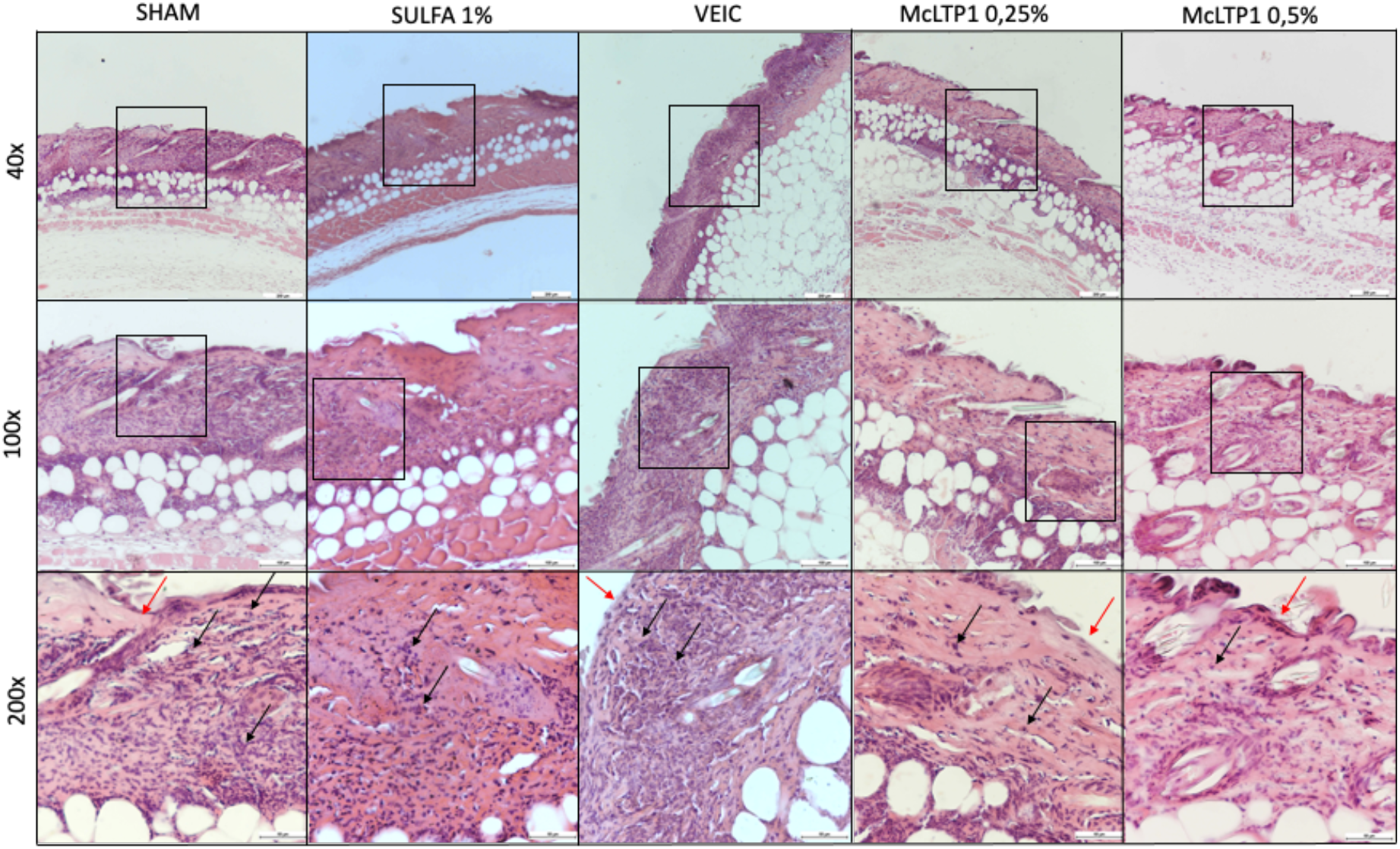
Photomicrographs of the ulcerated area on the skin after 3 days in the experimental groups (HE)

On the 7th day, there was a significant reduction in the scores evaluated in the Sulfa 1% and McLTP1 0.5% groups, in relation to Sham and Vehicle (Chart 1; Figure 3). Persistence of the inflammatory infiltrate was observed after the burn, however the presence of fibroblasts and fibrocytes was observed (Chart 1; Figure 3). McLTP1 0.5% was able to promote fibroblast proliferation and epithelium formation just below the wound crust, which was not observed in animals treated with 1% silver sulfadiazine. However, both treatments were effective in controlling the presence of inflammatory infiltrate. Furthermore, an increase in the presence of fibroblasts in the dermis was observed in the group treated with 0.5% McLTP1.

**Chart 1.** Representation by scores of the histological evaluation of burns

| **Groups**<br>**Time** | **SHAM** | **SULFA 1%** | **VEIC** | **McLTP1**<br>**0,25%** | **McLTP1**<br>**0,5%** |
| --- | --- | --- | --- | --- | --- |
| **3<sup>rd</sup> day** | 2 (1-3) | 2 (1-3) | 2 (2-3) | 2 (2-3) | 2 (1-3) |
| **7<sup>th</sup> day** | 3 (2-3) | 2(1-2) *<sup>δ</sup> | 2,5 (2-3) | 2 (1-2) | 1 (1-2) *<sup>δ</sup> |

**Figure 3.**
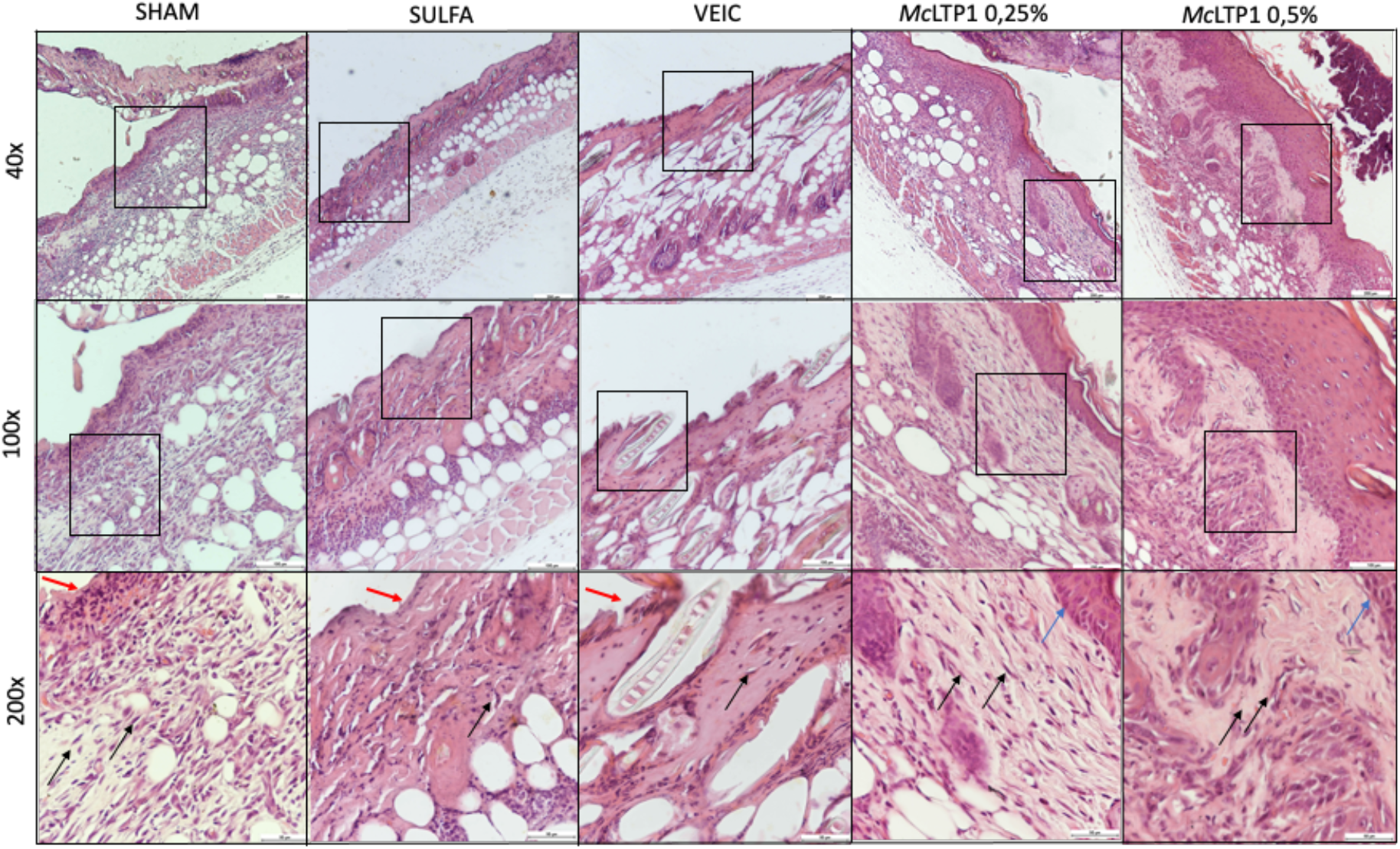
Photomicrographs of the ulcerated area on the skin after 7 days in the experimental groups (HE)

On the 14th day, it was observed that the animals treated with McLTP1 0.5% had a thicker epithelium compared to the other groups (Sham: 4.8 ±0.4; Sulfa 1%: 3.73 ±0.78; Vehicle: 4. 5 ±0.64; McLTP1 0.25%: 8.2 ±1.25 – Figure 4; 5).

**Figure 4.**
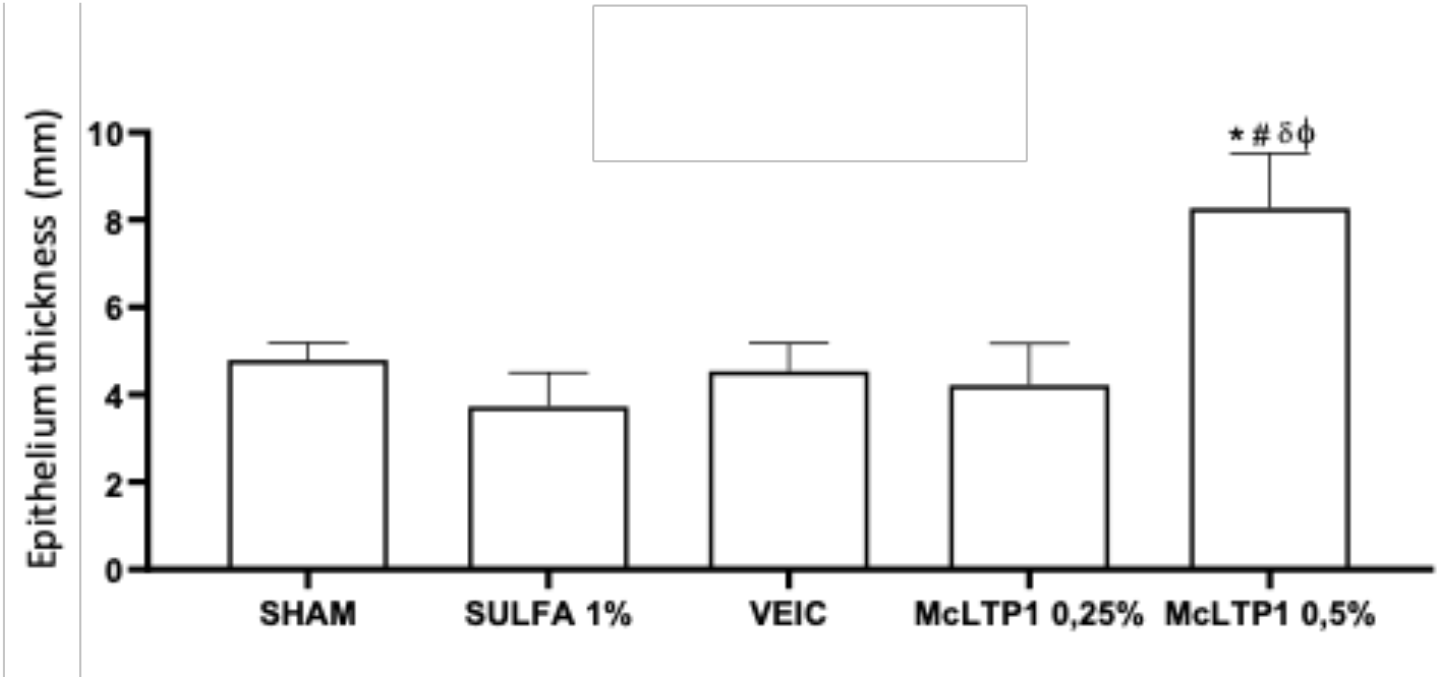
Effect of treatment with McLTP1 on the thickness of newly formed epithelium after 14 days.

**Figure 5.**
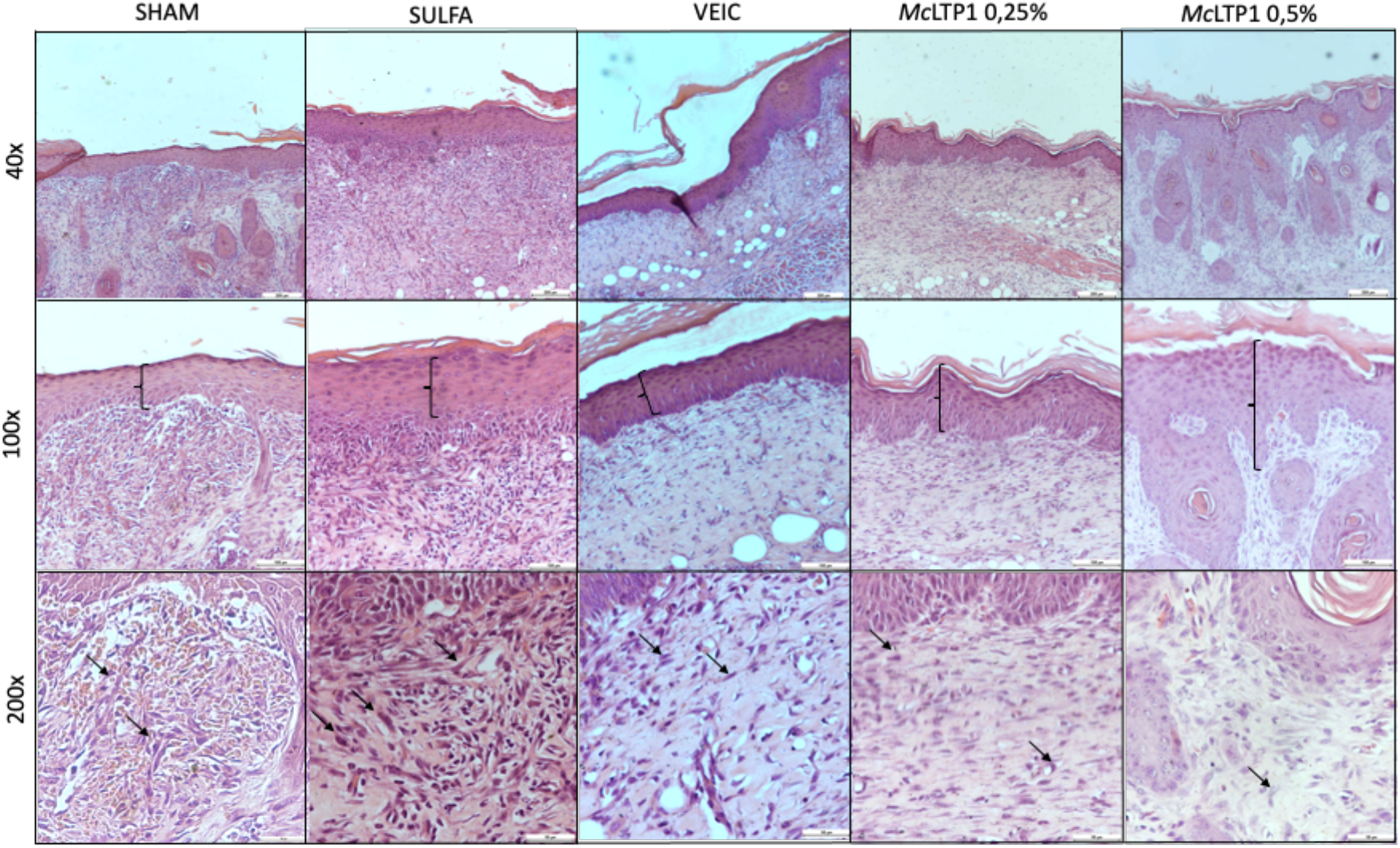
Photomicrographs of the ulcerated area on the skin after 14 days in the experimental groups (HE)

### McLTP1 reduced inflammation process after 3 days of superficial burns

Treatment with McLTP1 0.5%, when compared to the Sham group, significantly promoted the reduction of tissue levels of TNF-α, IL-1α and IL-6, increasing IL-10. On the other hand, treatment with McLTP1 at a concentration of 0.25% did not promote tissue alteration in the levels of any of the inflammatory mediators evaluated in this approach. 1% silver sulfadiazine reduced only TNF-α and IL-6 levels, not interfering with the tissue release of IL-1β and IL-10 levels.

**Figure 6.**
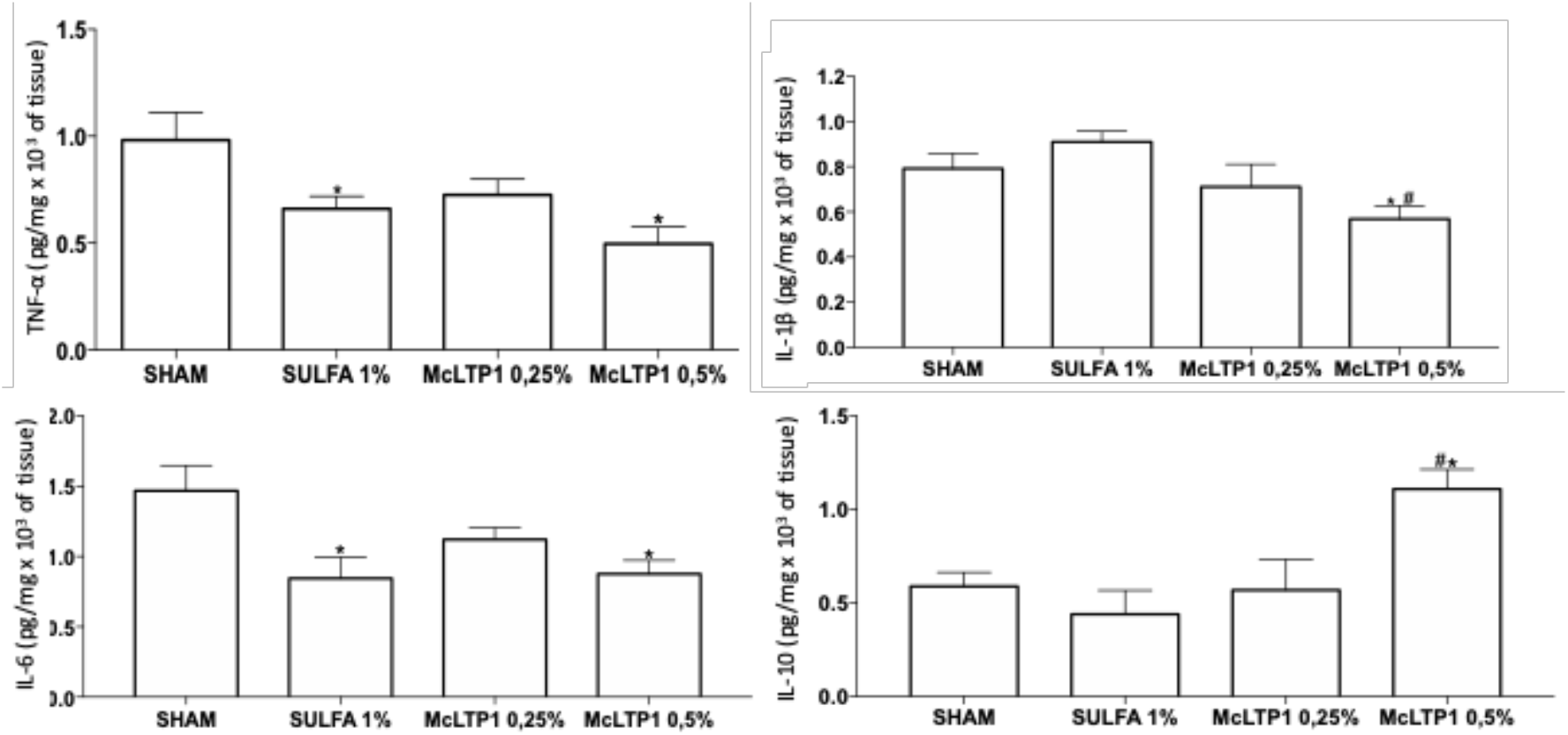
The levels of TNF-α, IL-1β, IL-6 and IL-10 were measured in the skin of the back submitted to superficial burn after 3 days of treatment. The results (picogram/mg 103 of tissue) were expressed as the mean ± standard error of the mean. ANOVA and Tukey’s post-test were used for comparisons between means. *p<0.05 represents a statistically significant difference in relation to the Sham group, #p<0.05 in relation to Sulfa 1% (N=5-6 animals/group).

### McLTP1 modulated angiogenesis and fibroblasts proliferation on superficial burns

On the 3rd day, McLTP1 at concentrations of 0.25% and 0.5% decreased VEGF levels compared to Sham, not differing from Sulfa 1%. On the 7th day, both concentrations of McLTP1 increased the levels of this growth factor in relation to Sham, and the increase in concentration of 0.5% was also significantly higher in relation to Sulfa 1%. On the 14th day, only the highest concentration of McLTP1 maintained a significant increase in levels of this mediator in relation to Sham. Considering the involvement of VEGF in angiogenesis, the presence of new vessels was evaluated macroscopically in photographs of the inner part of the skin on the back submitted to burns. The illustrations demonstrate the reduction of vessels in the animals in the groups treated with McLTP1 on the 3rd experimental day, followed by an increase in angiogenesis in these groups after 7 days. At the 14th day, the difference in the presence of new vessels between the Sham and McLTP1 0.5% groups is very evident, with an increase in the number and caliber of new vessels being seen in the experimental group compared to the control group.

On days 7 and 14 TGF-β presence was measured by ELISA. On the 7th day, Sulfa 1% and McLTP1 at concentrations of 0.25% and 0.5% increased TGF-β levels in relation to Sham. McLTP1 in its highest concentration was more effective in increasing TGF-β in relation to all groups. On the 14th day, only the Sulfa 1% group showed increased levels of TGF-β in relation to Sham, having decreased the levels of this growth factor in relation to the other groups.

**Figure 7.**
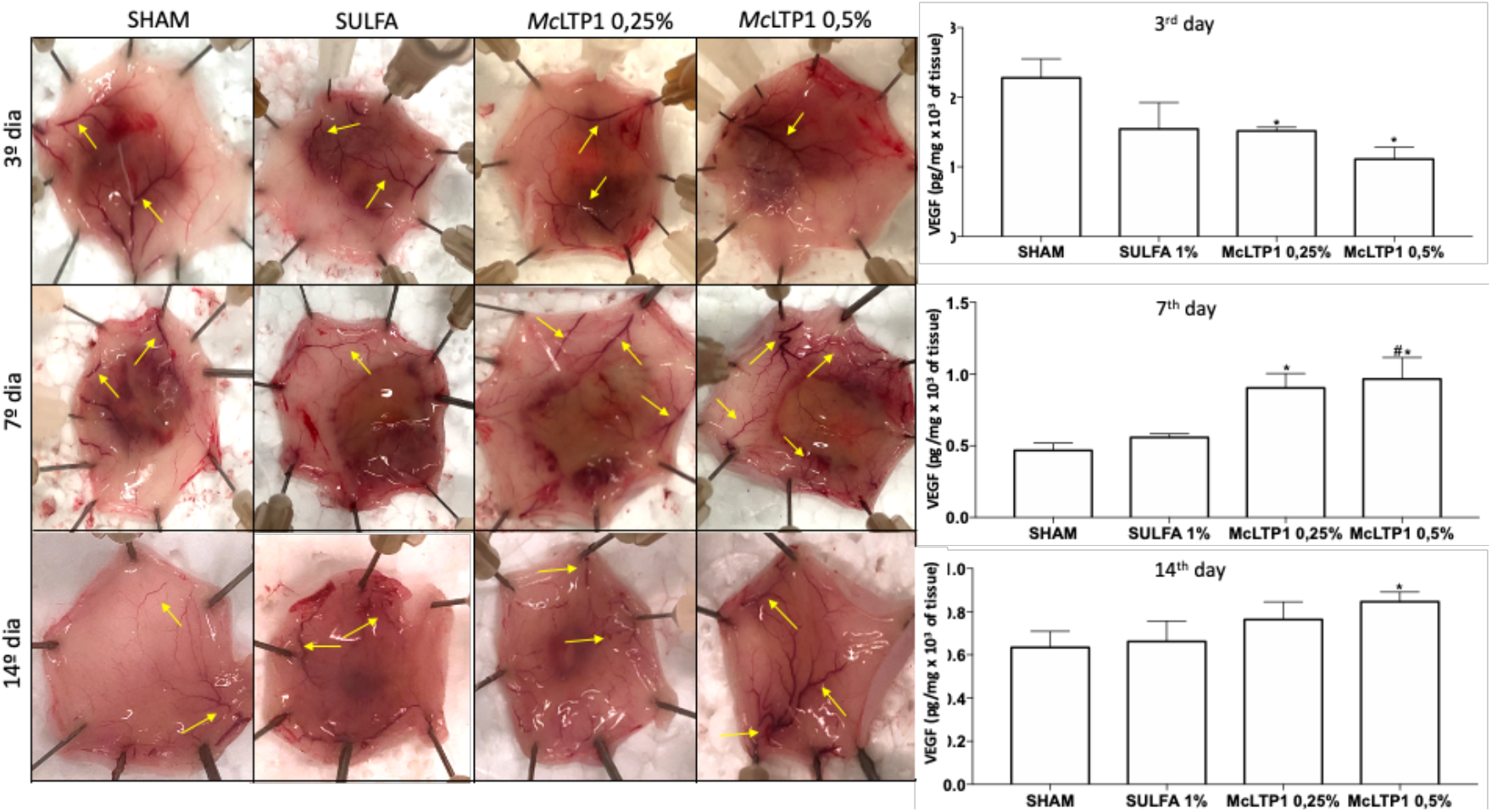
VEGF levels were measured in the skin on the back submitted to superficial burns after 3, 7 and 14 days of treatment. The results (picogram/mg 103 of tissue) were expressed as the mean ± standard error of the mean. ANOVA and Tukey’s post-test were used for comparisons between means. *p<0.05 represents a statistically significant difference in relation to the Sham group, #p<0.05 in relation to Sulfa 1% (N=5-6 animals/group). Yellow arrows indicate the new blood vessels.

**Figure 8.**
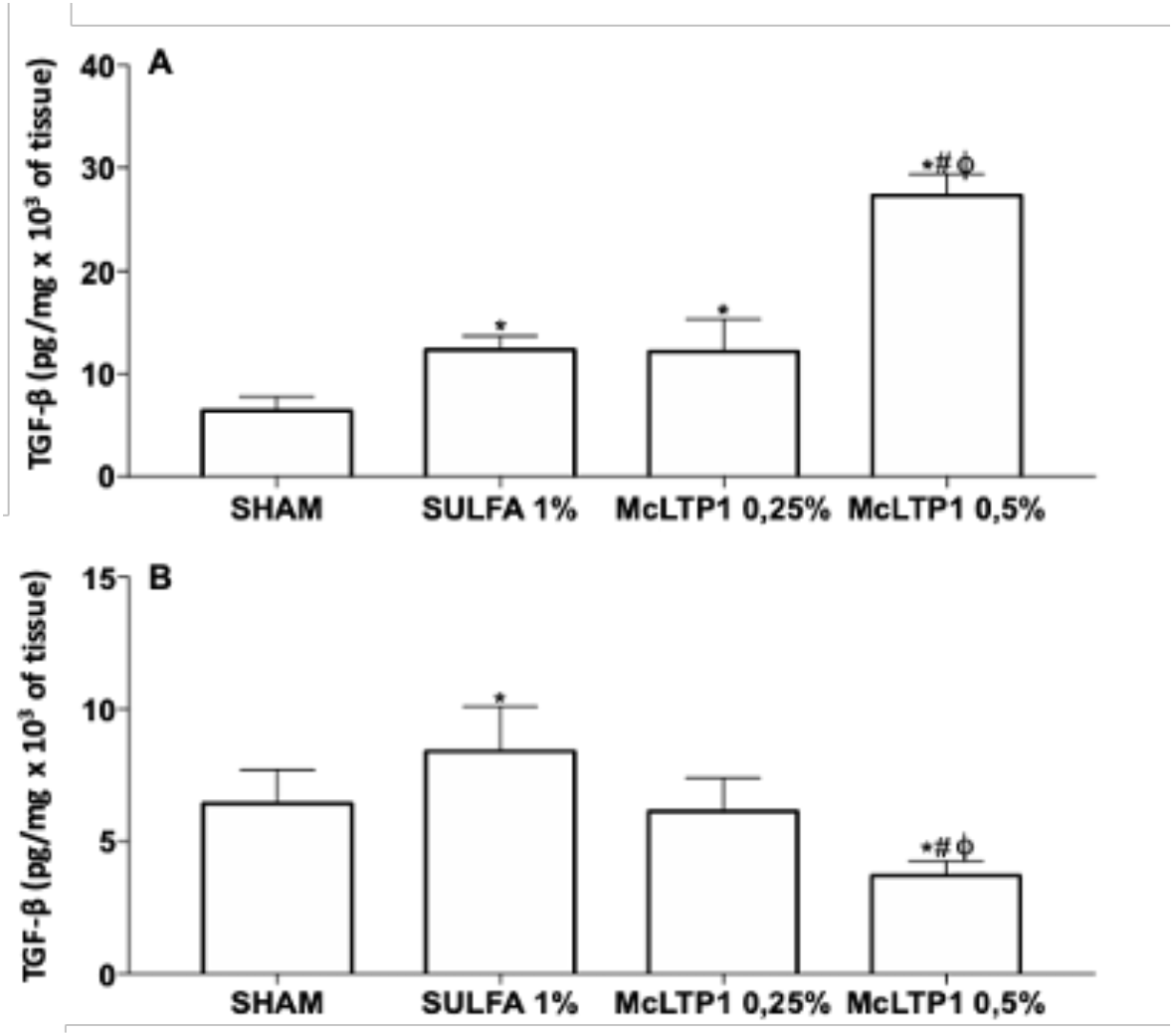
TGF-β levels (A and B) were measured in the skin on the back submitted to superficial burns after 7 and 14 days of treatment. The results (picogram/mg 103 of tissue) were expressed as the mean ± standard error of the mean. ANOVA and Tukey’s post-test were used for comparisons between means. *p<0.05 represents a statistically significant difference in relation to the Sham group, #p<0.05 in relation to Sulfa 1% and fp<0.05 in relation to McLTP1 0.25% (N=5-6 animals /group).

### McLTP1 reduced TNF-α expression after 3 days and increased FGF after 7 days of superficial burns

Skin samples collected on the 3rd day were processed for the evaluation of TNF-α expression by immunohistochemistry. The Sham group showed an increase in immunostaining for TNF-α while silver sulfadiazine 1% and McLTP1 0.5% decreased the staining of this cytokine in tissue samples.

**Figure 9.**
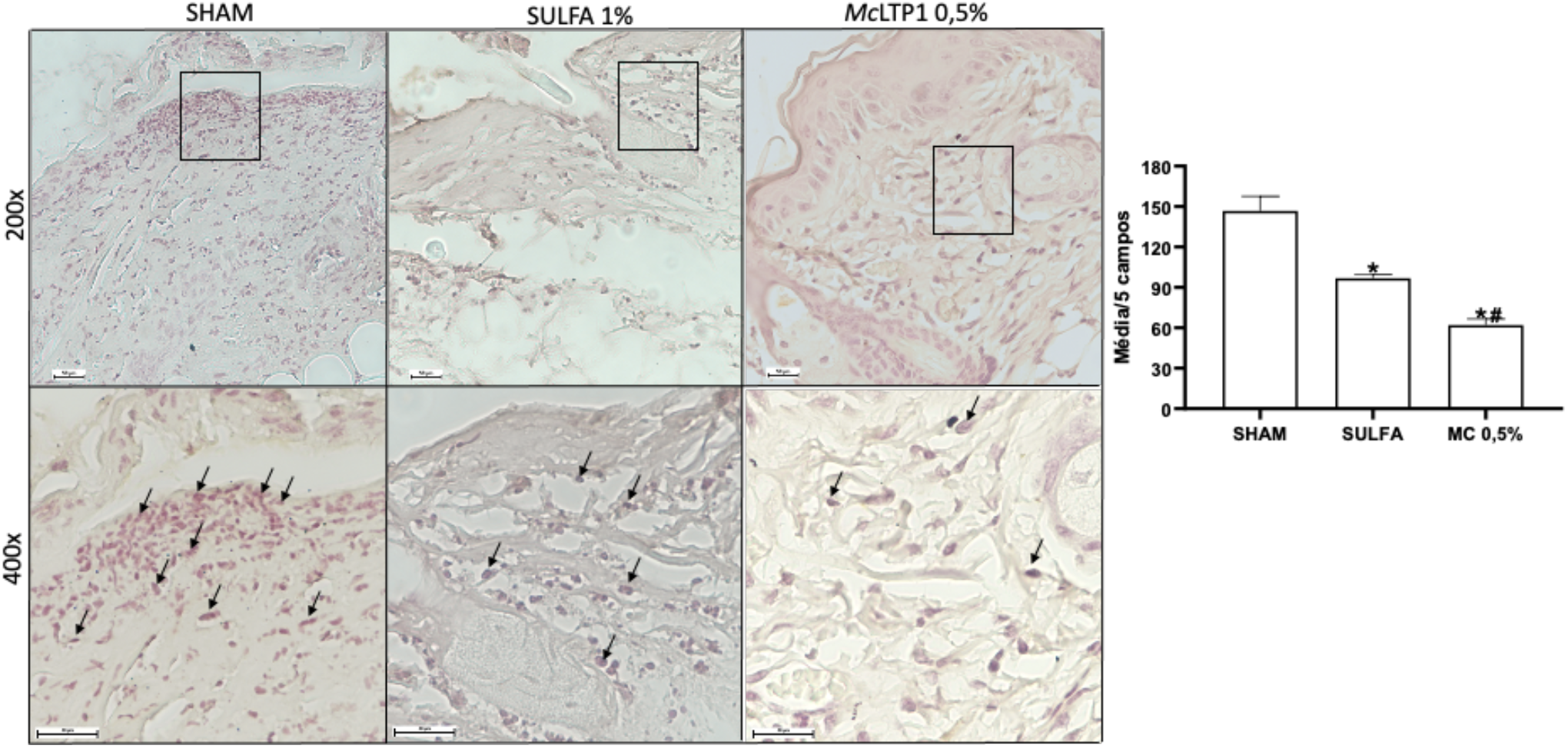
Quantification and photomicrographs of TNF-α immunostaining on the back skin of animals submitted to burns after 3 days

Skin samples collected on days 7 and 14 were targeted for immunohistochemistry for FGF. On the 7th day, there was a significant increase in FGF immunostaining in skin samples treated with McLTP1 0.5% compared to the Sham groups and Sulfa 1%. On the 14th day, the Sulfa 1% group demonstrated an increase in markings compared to Sham, while McLTP1 0.5% did not alter FGF expression in this period.

**Figure 10.**
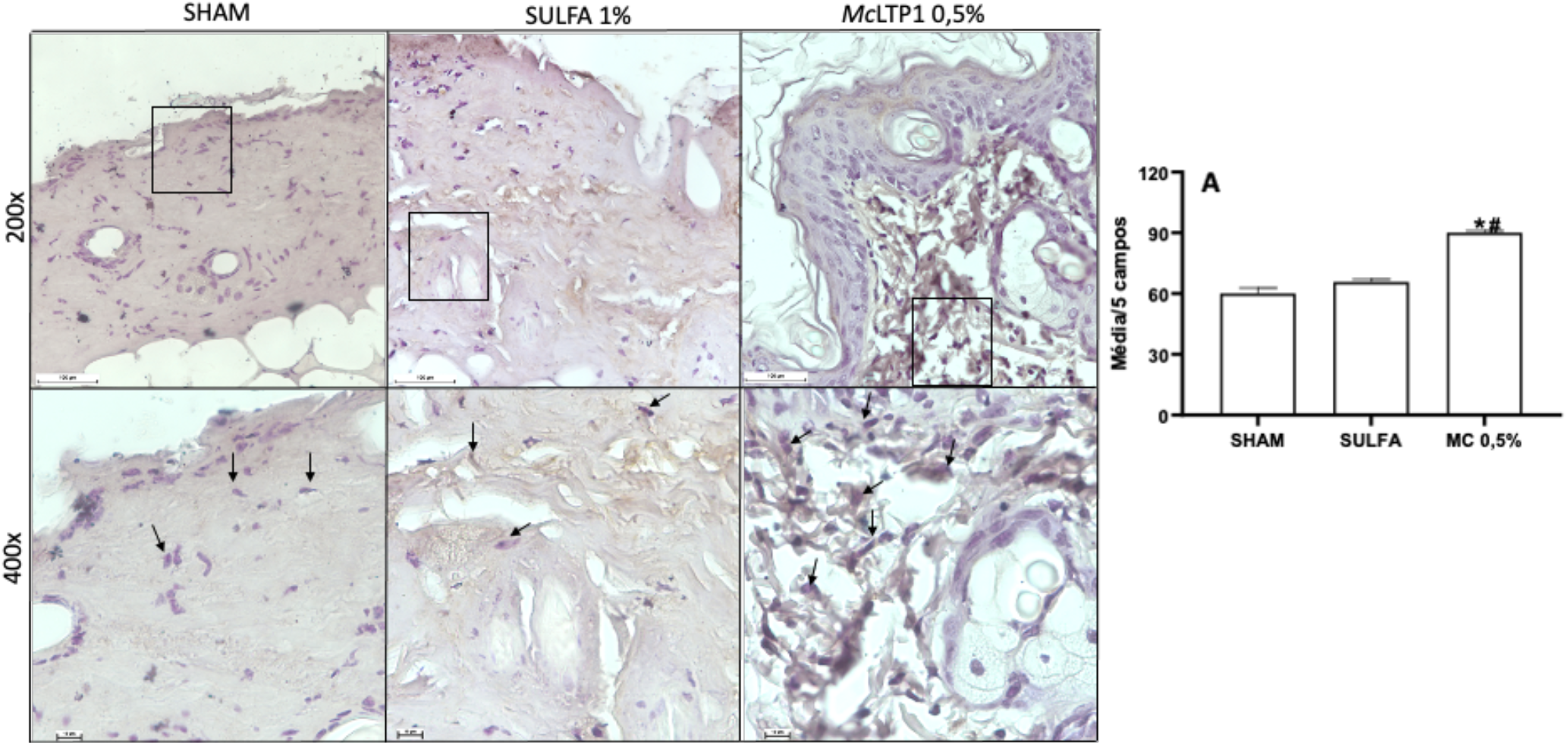
Quantification and photomicrographs of FGF immunostaining on the back skin of animals submitted to burns after 7 days

**Figure 11.**
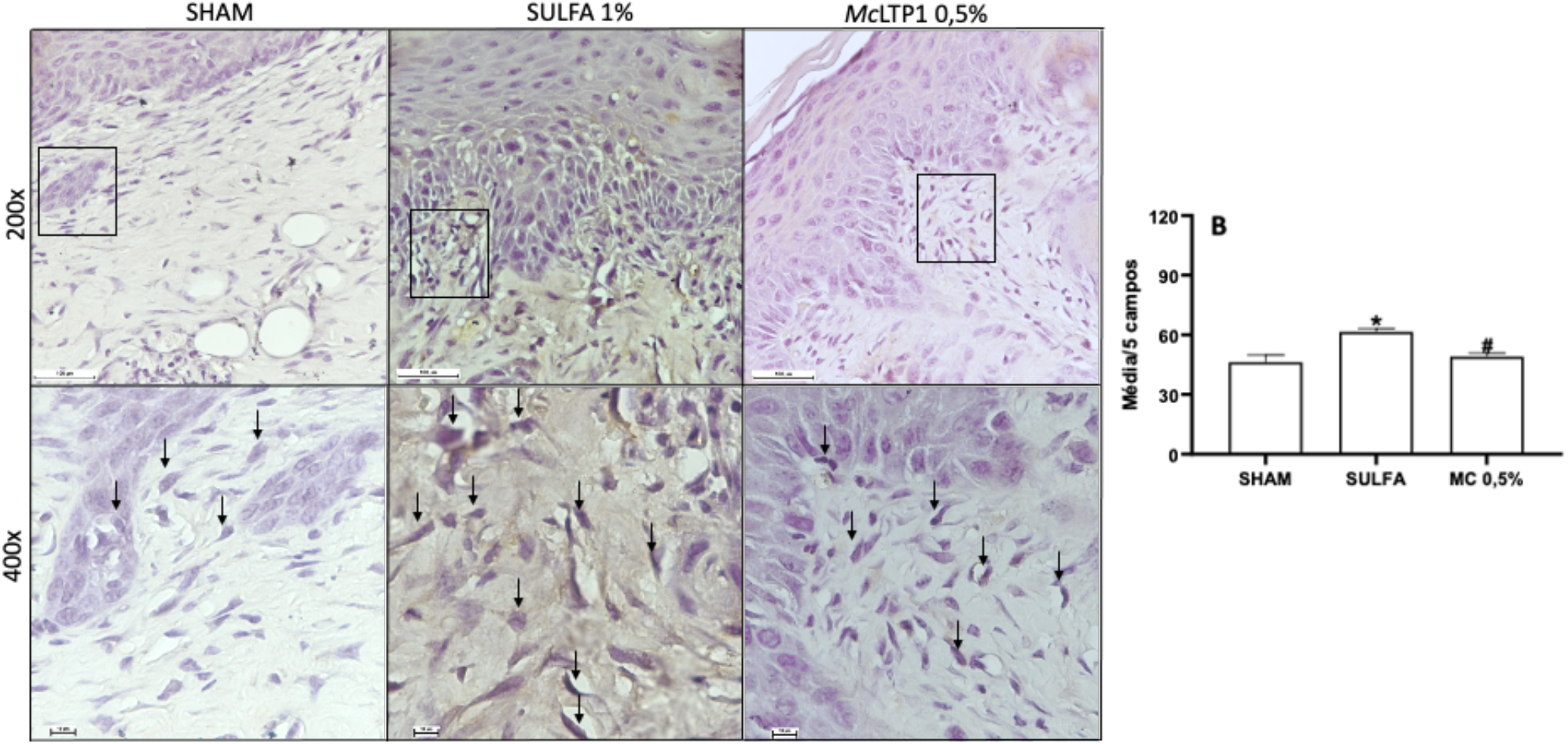
Quantification and photomicrographs of FGF immunostaining on the back skin of animals submitted to burns after 14 days

### *Mc*LTP1 reduced oxidative stress induced by burns

Skin samples collected on the 3rd day were processed to assess the levels of Treatment with McLTP1 at both concentrations significantly reduced MDA levels relative to Sham and Sulfa 1%. The conversion of NO2 to NO3 was decreased in groups treated with 1% silver sulfadiazine and 0.5% McLTP1. The McLTP1 protein, at both concentrations, promoted an antioxidant effect by raising GSH levels in relation to the Sham and Sulfa 1% groups at their two concentrations.

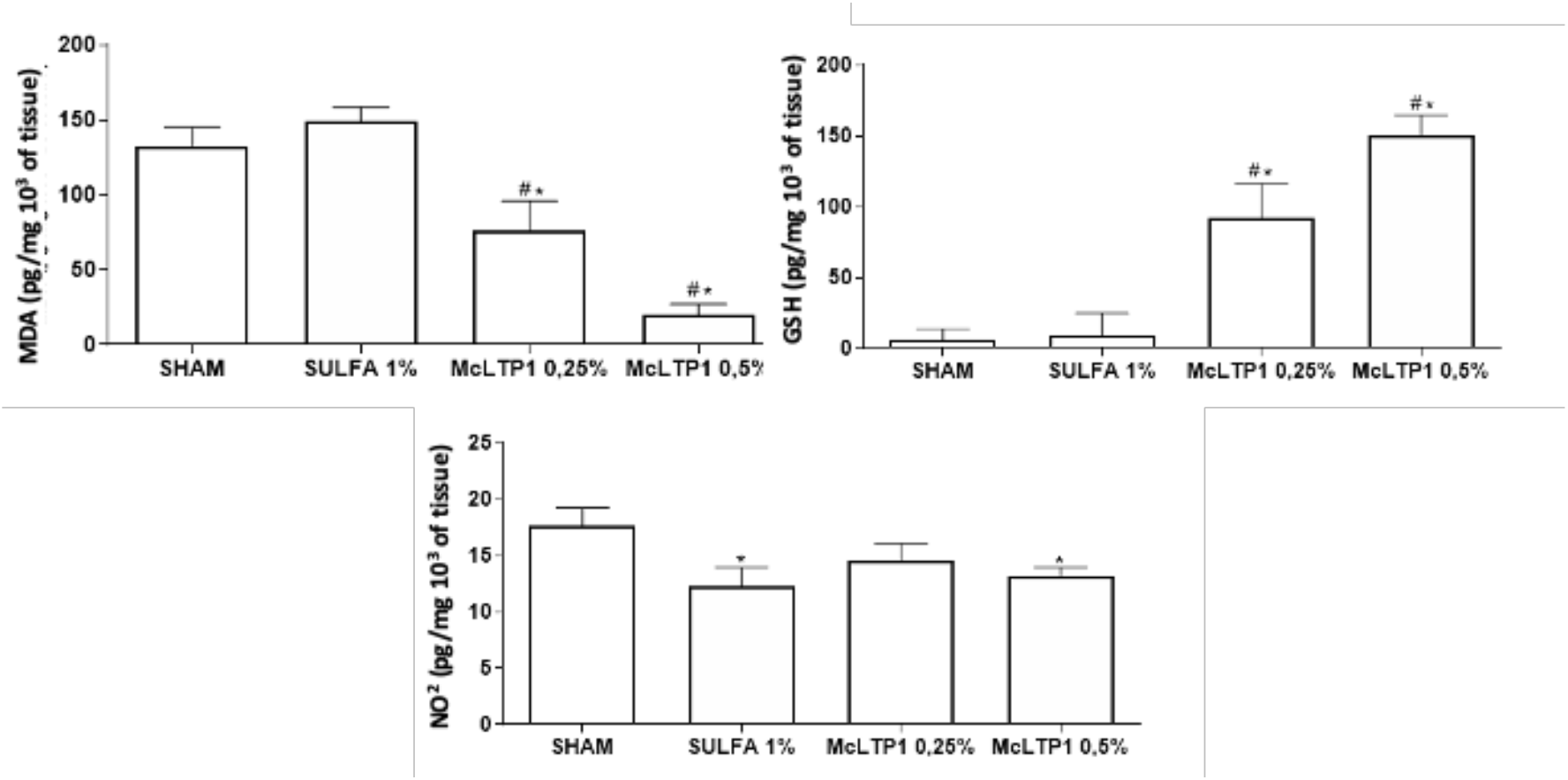
Oxidative stress was assessed by measuring MDA (A), GSH (B) and NO2/NO3 (C) in the skin on the back submitted to superficial burns after 3 days of treatment. The results (picogram/mg 103 of tissue) were expressed as the mean ± standard error of the mean. ANOVA and Tukey’s posttest were used for comparisons between means. *p<0.05 represents a statistically significant difference in relation to the Sham group, #p<0.05 in relation to Sulfa 1% (N=5-6 animals/group).

## Discussion

In this work, treatment with McLTP1 demonstrated a potent anti-inflammatory effect by reducing the inflammatory infiltrate, the activity of the enzyme myeloperoxidase, the levels of TNF-α, IL-1β, IL-6 and VEGF and increasing IL-10, as well as good antioxidant effect by decreasing malondialdehyde levels and NO conversion, as well as increasing reduced glutathione levels on the 3rd experimental day. Furthermore, it promoted epithelial repair by increasing epithelial thickness, by inducing VEGF, TGF-β and FGF after 7 days, and maintaining high angiogenesis after 14 days. Thus, this protein induced an excellent healing response, proven by the total closure of the wounds in the group treated with the 0.5% concentration.

Silver sulfadiazine 1% was chosen as positive control since it is recommended y healthy organization in Brazil^39^. However, its effect on burn wound healing does not seems to be effective when compared to other natural products as Aloe vera and bee’s honey^40,41^. This study also find that McLTP1 was more effective than silver sulfadiazine in repairing skins.

Superficial burns induced changes in tissue architecture of epithelium and dermis, presence of ulcers, intense inflammatory infiltrate in dermis and hypodermis after 3 days. Also induced increase on MPO activity, TNF-α, IL-1β and IL-6 and reduced IL-10 levels. This cytokines kinetics was also observed previously^42^. In the face of these findings, 3^rd^ day was chosen to evaluate inflammatory response. At de 7^th^ day, ulcers were still presents on histologic evaluation, but the inflammatory infiltrate seemed to be quickly replaced by fibroblasts and fibrocytes. So, this period was chosen to analyze transition of inflammatory to proliferate phase. On the 14^th^ day, epithelium was almost newformed, with high presence of fibroblasts and collagen fibers. Due to these alterations, the evaluation of epithelium repair was made at this time.

McLTP1 treatment did not altered scores at 3^rd^ day, even having reduced inflammatory cells, because ulcer presence elevated these scores. These results were similar to others in the same model^17^. Treatment with McLTP1 0.5%, as silver sulfadiazine, reduced scores of histologic evaluation only after 7 days of burn. The protein increased fibroblast and fibrocyte presence and epithelium repair, even in the presence of scabs. At 14^th^ day, the highest concentration of McLTP1 induced total closure of wounds with a thicker and more adhered epithelium when compared to the other groups.

Aiming to verify results of histologic analysis, the inflammatory profile of burn wounds and treatments with silver sulfadiazine and McLTP1 was evaluated. MPO activity, a marker of neutrophil presence, was increased after 3 days of burn, in according with other protocols^31^. Indeed, neutrophils are the more abundant cells after 48 hours of burns^43^. Beyond these cells, monocytes and macrophages also migrate to injury local and produce inflammatory cytokines, as TNF-α, IL-1β and IL-6^44^. Low levels of IL-10 after burn results to M1 macrophages that amplify Th1 lymphocytes response^45^. Only silver sulfadiazine and McLTP1 0.5% were capable of reduce MPO activity. Levels of TNF-α, IL-1β and IL-6 were reduced while IL-10 was increased by highest concentration of McLTP1. The modulation of cytokines was observed previously, in pep ulcer model^32^. Immunohistochemistry of TNF-α showed increased of immunostaining after 3 days, proving the importance of this mediator in early stages of burn injury^42–44^ and corroborating with our previous results. Silver sulfadiazine and McLTP1 reduced TNF-α expression, as expected from these anti-inflammatory effects.

Growing factors as VEGF and TGF-β are important in wounds healing since revascularization and fibroblasts recruitment are key steps to repairing process^46,47^. Angiogenesis was evaluated by VEGF levels analysis. This dosage was made at 3^rd^, 7^th^ and 14^th^ days, since a change in the revascularization pattern of the skin on the back of the animals was observed. VEGF participate on vascular changes of inflammation, so, it is expected to be in higher levels after burn injury^42^ what was observed in the study. So, the reduced levels of this factor after McLTP1 treatment show its anti-inflammatory effect, since inhibition of vascular activity may contribute to reduction of inflammatory cells chemotaxis, edema, and redness^48^.

VEGF dosage after 7 and 14 days showed different results, since McLTP1 increased the levels of this growing factor, demonstrating its repairing activity by inducing angiogenesis. VEGF presence was observed in high levels during all the stages of burn injury^42^.Potentialize vascular activity as seen after increase in VEGF levels on groups that received McLTP1 may be associated with faster healing. Other findings of literature also showed rapidly healing after increasing VEGF expression^49,50,51^.

At 7^th^ day, TGF-β levels increased after McLTP1 treatment, with the concentration of 0.5% being more effective than the other treatments. On the 14^th^ day, McLTP1 0.5% reduced TGF-β levels in the same way. The upregulation of this factor after 7 days may be related to wound healing acceleration, since it was demonstrated that burns repair also shows increasing of TGF-β in 7 days^42^. The downregulation observed on the 14^th^ day can be attributed to the fact that exacerbated increase of TGF-β causes cicatricial hypertrophy and keloids, resulting in malformed scars^52^. So, controlling TGF-β levels by McLTP1 demonstrate its potential to performs a wound healing with better quality.

In order to confirm findings of TGF-β levels, immunostaining of FGF, a growing factor for fibroblasts^44^ was taken. FGF participation on burn wound healing by natural products was previously related, with high effectiveness^53,54^. In this study, McTLP1 increased FGF expression after 7 days, in accordance with TGF-β levels and literature previous relates^53–54^. At 14^th^ day, silver sulfadiazine 1% treatment resulted in increased FGF immunostaining at late stage of burn wound healing, but treatment with McLTP1 do not alter FGF expression. However, it was observed a complete wound closure at this period. Together, these findings suggests that cellular modulation of McLTP1 be more intense at 7^th^ day, during transition phase of healing.

Oxidative stress was evaluated at 3^rd^ day, since it has a bidirectional relation with inflammatory response mediated by neutrophils and increase on cellular and tissue oxidative damage^55,6^. Studies related that malondialdehyde (MDA) is a marker of lipid peroxidation^56,57,58^ so its levels were analyzed after burns. Thermal injury increases MDA levels which were reduced by both concentrations of McLTP1. This result is in accordance with other protocols^32^. Metabolization of nitric oxide (NO) in NO2 and NO3 also contributes to increase oxidative stress and tissue damage^59^. Indeed, superficial burn enhanced NO2/NO3 levels, while McLTP1 0.5% and silver sulfadiazine 1% prevent NO metabolization. Reduced glutathione (GSH) is a tripeptide that acts in tissue and cellular protection in the presence of oxidative stress, promoting an antioxidant effect^56,60,58^. Only McLTP1 increased GSH levels after burn, showing its potential to reverse injuries caused by superficial burns. Similar results were observed previously^32^.

In short, McLTP1 demonstrated an excellent healing response in stimulating wound contraction and promoting total wound closure in the group treated with the 0.5% concentration. These findings were confirmed by the histological evaluation using scores and by measuring the epithelium, where McLTP1 0.5% reduced the inflammatory infiltrate and promoted greater re-epithelialization. It is suggested that these effects were mediated by the control of the inflammatory response, as well as by the up regulation of growth factors, as assessed by MPO measurement, ELISA and immunohistochemical staining. Additionally, these effects seem to have been enhanced by its antioxidant action.

## Conclusion

The lipid transfer protein isolated from Noni seeds (McLTP1) demonstrated healing potential in an experimental model of superficial burns in mice by modulating the inflammatory response and oxidative stress, and accelerating tissue repair. It is suggested that McLTP1 may represent an interesting pharmaceutical approach for the treatment of wounds in general.

## Acknowledgments

This work was supported by research grants from Fundação Cearense de Apoio ao Desenvolvimento Científico e Tecnológico (FUNCAP), Conselho Nacional de Desenvolvimento Científico e Tecnológico (CNPq) (Award Number: SPU 07971628/2020 in called SUS/PPSUS - CE 02/2020), and scholarship from Coordenação de Aperfeiçoamento de Pessoal de Nivel Superior (CAPES).

